# No evidence for transient transformation via pollen magnetofection in several monocot species

**DOI:** 10.1101/2020.05.01.071266

**Authors:** Zuzana Vejlupkova, Cedar Warman, Rita Sharma, Henrik Vibe Scheller, Jenny C. Mortimer, John E. Fowler

## Abstract

The development of rapid and efficient transformation methods for many plant species remains an obstacle in both the basic and applied plant sciences. A novel method described by Zhao et al. (2017) used magnetic nanoparticles to deliver DNA into pollen grains of several dicot species, and one monocot (lily), to achieve transformation (“pollen magnetofection”). Using the published protocol, extensive trials by two independent research groups showed no indication of transient transformation success with pollen from two monocots, maize and sorghum. To further address the feasibility of magnetofection, lily pollen was used for side-by-side trials of magnetofection with a proven methodology for transient transformation, biolistics. Using a Green Fluorescent Protein reporter plasmid, transformation efficiency with the biolistic approach averaged 0.7% over three trials. However, the same plasmid produced no recognizable transformants via magnetofection, despite screening >3500 individual pollen grains. We conclude that pollen magnetofection is not effective for transient transformation of pollen for at least three species of monocots, and suggest that efforts to replicate the magnetofection protocol in dicot species would be useful to fully assess its potential.

ARISING FROM Zhao et al. *Nature Plants* https://doi.org/10.1038/s41477-017-0063-z (2017)

---

Except for a few species, achieving stable transformation of plants remains a major hurdle, hindering efficient progress towards both basic and applied scientific goals. Thus, the novel approach of pollen magnetofection, described by Zhao et al. ^1^ was greeted as a potentially transformative technology. In this approach, magnetic nanoparticles deliver exogenous DNA into pollen grains, which then generate genetically modified progeny incorporating that DNA into their genome. Independently, our two groups set up trials to investigate the effectiveness of this approach in two monocot species, maize and sorghum. Both species, which require specialized expertise and demanding methodologies to achieve transformation ^2^, are of interest as models for basic biological investigation, as well as for their agricultural impact. Moreover, both species produce readily accessible pollen, raising the possibility that the pollen magnetofection approach could be easily implementable with these grasses.

Zhao et al. focused primarily on dicot species (cotton, pepper, pumpkin, and cocozelle). However, although stable transformation was not achieved, some success was reported for one monocot, lily (*Lilium brownii*). With lily, transient transformation of pollen was reported at ∼90% efficiency, as detected by staining for β-glucuronidase (GUS) activity following magnetofection with a *GUS* reporter plasmid. Thus, as a prelude to attempting stable transformation in sorghum and maize, we chose to focus on testing transient transformation efficiency in pollen, as results could be assessed rapidly (typically within a day). However, initial experiments with sorghum indicated that *GUS* reporters would not be ideal, due to positive GUS staining in control, non-transformed pollen grains (Supplemental Figure 1). Other published reports also indicate that GUS activity is detectable in the male gametophyte of a number of plant species ^3,4^, spurring our labs to use green fluorescent protein (GFP)-based reporter plasmids as an alternative. For these experiments, we chose plasmids with promoters known to be highly expressed in grass pollen (maize *Zm13*, rice *Actin1*, or maize *ZmUbiquitin1*). However, following the published protocol, over twenty magnetofection trials with either sorghum or maize pollen generated no indication of plasmid-induced transient expression of fluorescence (Supplemental Figure 2, Supplemental Table 1) (over 50,000 pollen grains screened).

To determine whether our lack of successful transient transformation was due to a grass-specific resistance to magnetofection, we turned to lily pollen as a model. Using lily pollen afforded us the ability to compare transformation efficiency via magnetofection vs. biolistics, a well-established methodology ^5^. The magnetofection protocol is simple enough to allow side-by-side treatment alongside the biolistic protocol, enabling assessment of transformation efficiency in subsamples of the same population of pollen (see Supplemental Methods). Briefly, pollen was collected from cut lily flowers (*Lilium* var. Santander) by vortexing anthers in pollen germination media (PGM); biolistic microcarriers and magnetic nanoparticles were coated with the *Zm13* promoter-driven GFP reporter plasmid; and nanoparticles or microcarriers were delivered to pollen via exposure to a magnet or biolistic bombardment, respectively. Treated pollen samples were then incubated overnight in PGM, and imaged to detect reporter expression the following day via fluorescence microscopy. Transformation efficiency was determined via blind assessment of the number of transformed pollen grains/tubes in >10 randomly chosen microscopic fields for each replicate. Over 1,000 pollen grains or tubes were screened for each experimental treatment in each of three trials (Supplemental Table 2). We followed the Zhao et al. protocol as published, with one exception: we used a PGM formulation ^6^ that, in our hands, gave improved pollen germination frequencies relative to that given by Zhao et al.

Transformed pollen grains and tubes were clearly recognizable by the expression of GFP fluorescence above background following biolistic bombardment with the reporter plasmid (Fig. 1). Biolistic transformation efficiency ranged from 0.4 to 1.1% in the three trials, averaging 0.7% (Table 1). As expected, no transformed pollen was found in the negative control biolistic treatments, i.e., samples bombarded with microcarriers lacking plasmid DNA. However, we also observed no transformed lily pollen in the magnetofection treatments, despite screening nearly 4,000 pollen grains (Table 1). Using the Cochran-Mantel-Haenszel test to assess the repeated trials indicates a statistically significant difference between the positive control biolistic method and both the negative control and the magnetofection protocol (p<10^−6^ for both).

**Figure 1.**
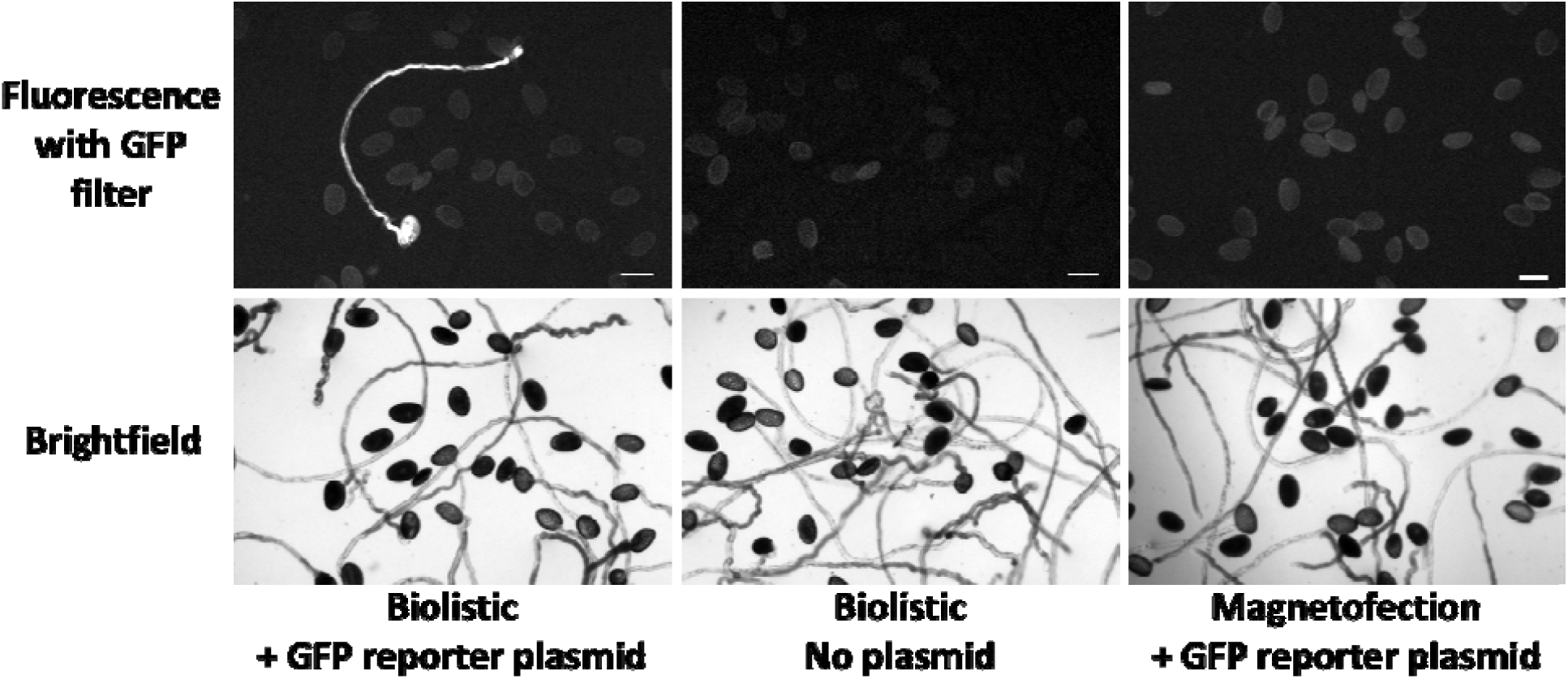
Strong GFP fluorescence is detectable in lily pollen and pollen tubes following biolistic bombardment with the *Zm13 promoter::*GFP reporter plasmid, whereas no green fluorescence is detected above background following magnetofection with the same plasmid. Scale bar = 100 µm.

**Table 1.** Summary of side-by-side magnetofection and biolistic transient transformation efficiency with lily pollen, 3 trials

|  | **Biolistic,**<br>**+ plasmid**<br>**DNA** | **Biolistic,**<br>**no DNA** | **Magnetofection,**<br>**+ plasmid DNA** |
| --- | --- | --- | --- |
| Total transformants (pollen grains & pollen tubes) | 33 | 0 | 0 |
| Total pollen in imaged fields | 4651 | 5559 | 3731 |
| Transformation efficiency | 0.71% | 0 | 0 |
| p value, Cochran-Mantel-Haenszel test (vs. biolistic positive control) |  | 4.09E-09 | 5.02E-07 |

In conclusion, using the protocol described by Zhao et al., we were unable to reproduce any evidence of transient transformation in lily pollen via magnetofection. Although we cannot rule out transient transformation at very rare frequencies, we suspect that the report of ∼90% transient expression efficiency following magnetofection of lily pollen was due to endogenous GUS activity, rather than that from the *GUS* reporter used for assessing magnetofection success. Finally, given our lack of success in lily, sorghum and maize despite extensive trials, we also could not generate evidence to support the idea of broad applicability for pollen magnetofection across monocots. We note that we cannot address the utility of pollen magnetofection for stable transformation, as we have not tested for genomic integration of reporter DNA. However, given our results, we believe it is important for groups working in dicot species, particularly those cited as successfully transformed (cotton, pepper, and pumpkin), to provide data regarding attempts to replicate both transient and stable transformation success via pollen magnetofection.

## Supplemental Material

**Supplemental Figure 1**. Sorghum pollen exhibits GUS activity in the absence of a GUS reporter plasmid. Scale bar = 100 µm.

**Supplemental Figure 2**. No fluorescence above background was detected in sorghum pollen following magnetofection with the *pZmUbi1*-GFP reporter plasmid. Scale bar = 100 µm.

## Supplemental Methods

**Supplemental Table 1**. Magnetofection trials and parameters tested with sorghum and maize pollen, assessed for transient transformation by screening for GFP reporter expression.

**Supplemental Table 2**. Lily pollen transformation efficiency via biolistics vs. magnetofection, details of three trials.

## Acknowledgments/funding

R.S. acknowledges IUSSTF-DBT for GETin fellowship. This work was funded in part by NSF grant IOS-1832186 (ZV, CW, JEF), and by DOE Joint BioEnergy Institute (http://www.jbei.org) supported by the U.S. Department of Energy, Office of Science, Office of Biological and Environmental Research through Contract DEAC0205CH11231 between Lawrence Berkeley National Laboratory and the U.S. Department of Energy (RS, HVS, JCM).

