## Supplemental_Figures_and_Methods for "No evidence for transient transformation via pollen magnetofection in several monocot species"

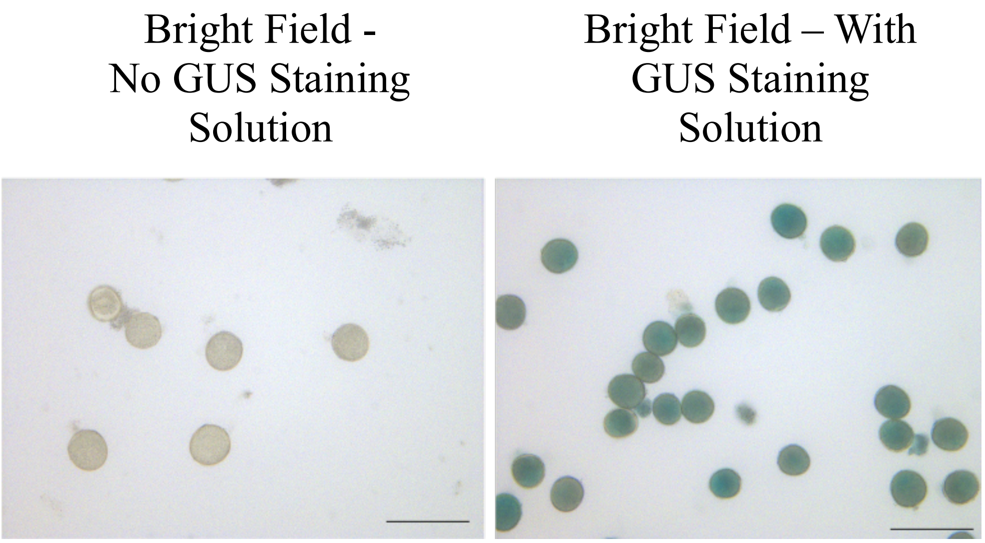


**Supplementary Figure 1.** Sorghum pollen exhibits GUS activity in the absence of a GUS reporter plasmid. Scale bar = 100 µm.

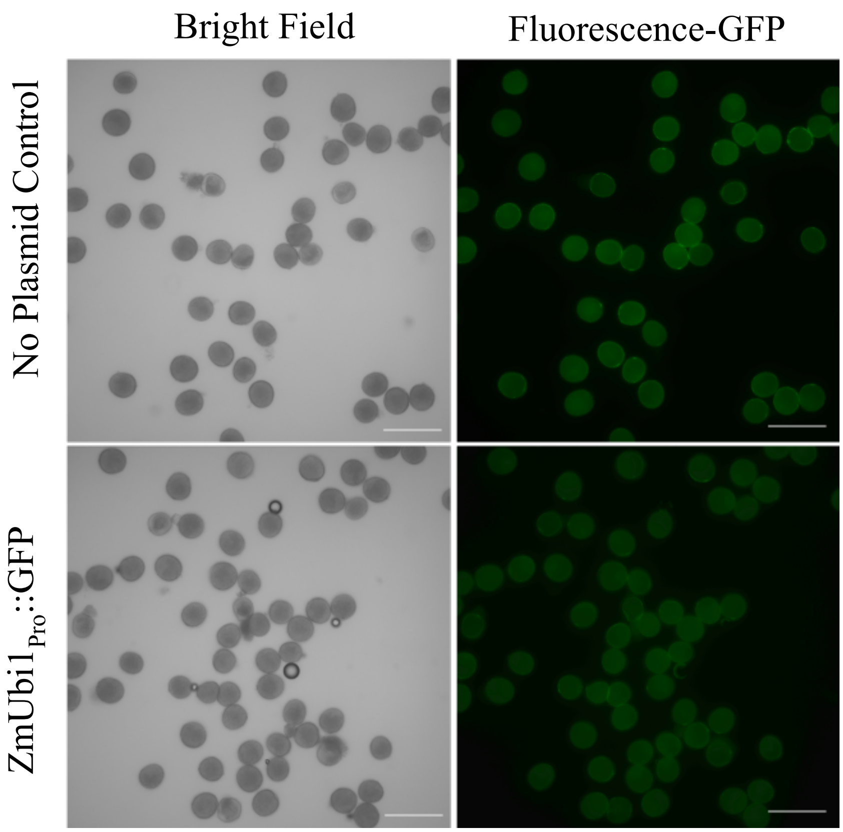


**Supplementary Figure 2.** No fluorescence above background was detected in sorghum pollen following magnetofection with the pZmUbi1-GFP reporter plasmid. Scale bar = 100 µm.

### Supplemental Methods

#### Sorghum Pollen Magnetofection Trials

*Sorghum bicolor* (L.) Moench cv Tx430 plants were grown in the University of California Berkeley greenhouse (maintained at 21 ºC minimum at night and 26-34 ºC during the day, with a 16 h photoperiod and supplemental light of 200 μmol $m^{-2}$ $s^{-1}$). Sorghum pollen was collected within hours after anthesis, and the method of Zhao et al. was followed [1]. Freshly collected pollen was directly shed in a medium containing 0.7 M sucrose, 3 mM calcium nitrate and 2.43 mM boric acid, suspended by gentle mixing, and an aliquot of 300 μl was added to a well of a 24-well plate. The plate was centrifuged for 2 min at 50 *x*g so that all the pollen collected at the base of the plate. Plasmid DNA (1 μg, C286-$Ubi_{PRO}$::GFP [2]) was mixed with Polymag beads at a 1:1 mass ratio, and incubated at 21 ºC for 15 min. The pollen-containing plate was placed upon the MagnetoFACTOR-24 plate (Chemicell), the Polymag/DNA complex was added to each well, and it was incubated for 30 minutes. The plate was then incubated at 21 ºC in the dark for about 24 hours without the magnetoFACTOR plate. Pollen was observed using a Leica D4000B fluorescence microscope coupled with a Leica DC500 camera and a GFP filter. For the GUS assay, sorghum pollen was incubated in GUS staining solution (0.5 mg/ml X-Gluc in dimethylformamide, 0.05% Triton X-100 and 50 mM sodium phosphate buffer, pH 7.0) or water (for control) for about 6 hours.

### Maize Pollen Magnetofection Trials

*Zea mays* plants were grown in the Oregon State University greenhouse under standard conditions (16 hrs light, 8 hrs dark; 27 °C day/21 °C night). Maize pollen was collected at anthesis and subjected to magnetofection within one hour after collection, following the method of Zhao et al. [1], with additional variations in relevant parameters documented in Supplemental Table 1. For the magnetic nanoparticle/DNA complexes, plasmid DNA from one of two GFP reporter constructs (pUC19-260Zm13::GFP or pBS_Act1promTSS::mGFPER) was mixed with Polymag beads (Oz Biosciences) at recommended ratios, and incubated at RT for 30 min. Pollen-containing media was mixed with recommended amounts of Polymag/DNA complex, and placed upon the MF10000 Super Magnetic Plate (Oz Biosciences), varying exposure parameters as detailed in Supplemental Table 1. The samples were incubated at RT in the dark following exposure to the magnet, and assessed for GFP expression using a Zeiss Axiovert S100 microscope equipped with an HBO 50 Illuminator, GFP filter and Qimaging (Retiga ExI) camera, at times specified in Supplemental Table 1.

**Protocol:**

**Biolistic transient transformation of lily pollen, side-by-side with magnetofection**

Adapted from [1, 3, 4]

The biolistic transformation [3,4] can be performed side-by-side with the magnetofection procedure [1], in order to use the same batch of fresh pollen for both. This procedure describes biolistic transformation, but also refers to steps of the magnetofection when timing is critical.

Gold microcarrier particles (1.0 µm; Bio-Rad, cat no. 165-2263) can be prepared ahead of time and stored in aliquots at -20 °C for up to 1 year. To prepare gold microcarriers, first wash with ethanol and water, then resuspended in 50 % sterile glycerol to 60 mg/ml.

**PROCEDURE**

**Coat gold particles with plasmid DNA**

1) Vortex 25 µl aliquot of gold particles in 1.5 ml Eppendorf tube for 5 min.

2) First, add 10 µl of 0.1 M spermidine (Sigma-Aldrich cat. No. S-2626). Second, add 5 µl of plasmid DNA (1µg/µl) and keep vortexing for another 1 min. Third, slowly add 25 µl of 2.5 M $CaCl_2$ solution while mixing. Keep vortexing for 3 min.

3) Centrifuge the microcarrier solution in benchtop centrifuge briefly for 5 s and remove the supernatant.

4) Wash with 70 µl of fresh 70% ethanol without disturbing the pellet. Remove 70% ethanol and replace with 70 µl absolute ethanol.

5) Remove the ethanol carefully without disturbing the pellet and resuspend gold particles/pellet with 24 µl of absolute ethanol. Aliquot 6 µl of particle suspension onto each microcarrier disc and let them air-dry.

Note: It is essential to completely resuspend the gold particles by pipetting up and down and by gentle vortexing. Make sure that the particles are evenly distributed in the center of the macrocarriers.

· ***Magnetofection step 1*:** While the macrocarriers are drying, mix together MNP (Polymag transfection reagent, Oz Biosciences) and DNA plasmid to form MNP-DNA complexes and let incubate for 30 min at RT.

**Preparation of pollen grains for transformation**

6) Connect filtering flask fitted with Buchner funnel into the vacuum system (water system or pump).

7) Harvest 8 – 10 anthers from 4 – 5 lily flowers (Oriental lily var. Santander) and place them into 20 ml of pollen germination medium (PGM) in 50 ml Falcon tube. Vortex vigorously for 1 min to release the pollen grains into the medium and remove the anthers with forceps.

· ***Magnetofection step 2:*** Remove 4 x 1ml aliquots of pollen suspension for magnetofection. Each 1 ml aliquot is mixed with MNP-DNA complexes, then transferred into a well of a tissue culture plate. Place plate on the magnet (MF10000 Super Magnetic Plate, Oz Biosciences), and proceed with 30 min magnetofection as described (Zhao et al 2017).

8) Remove a 1.5 ml aliquot of pollen suspension for a pollen germination control (untreated pollen) and plate in a glass bottom cell culture dish as described below.

9) Pre-wet 70 mm filter paper (Whatman #1) with pollen germination medium, place into the Buchner funnel, with vacuum on low.

10) Vacuum-filter the remaining pollen suspension onto the pre-wetted filter paper to collect pollen grains evenly distributed on top of the filter paper.

11) Transfer the filter paper (pollen side up) into a new 100 mm plastic Petri plate.

**Transformation**

12) Set up the PSD-1000/He particle delivery system (Bio-Rad, cat no. 165-2257) as follows: Helium pressure: 1,100 PSI; Target distance 6 cm; Chamber vacuum 26 inHg; Gap width 3/8".

13) Bombard pollen grains on the filter paper three times at three different positions (turn the plate ~120˚ in the plate holder between each shot), recommended by Wang & Jiang 2011 to increase the transformation efficiency.

14) Immediately after bombardment cut off the outer edge of the filter paper (~ 1.5 cm) and discard. Wash pollen from the remaining center filter disc in a 50 ml Falcon tube with 5 ml PGM.

15) Plate several dilutions of washed pollen in PGM into glass bottom cell culture dishes (Electron Microscopy Sciences Cat no. 70674-02). Remove excess PGM, leaving only a thin layer of pollen-PGM suspension.

16) Incubate at RT in the dark overnight.

· ***Magnetofection step 3:*** After 30 min of magnetofection, plate and incubate magnetofected pollen-PGM suspension as described above.

17) As a negative control, proceed with harvesting another 8 – 10 anthers from 4 – 5 of the same flowers as above and follow the same protocol for pollen collection, filtration, bombardment and plating as above. This bombardment is with microcarriers that have not been coated with the plasmid DNA. A separate batch of pollen is used for the negative control, as it would be infeasible to perform the side-by-side positive control bombardment and magnetofection experiments in a timely manner.

**Imaging**

All imaging was done the day following transformation using a Zeiss Axiovert S100 microscope equipped with an HBO 50 Illuminator, GFP filter and Qimaging (Retiga ExI) camera. For each treatment, at least 10 randomly selected, non-overlapping fields of view were imaged with a 4X objective. Pollen in each field of view was imaged with both transmitted light (6 ms exposure time) as well as a GFP filter set (Chroma #41017 - 470/40 excitation, 495 long-pass dichroic, 525/50 emission) (800 ms exposure time) using μManager 2.0.0-gamma1 20190730 software [5].

**Quantification of Transformation Efficiency**

Fluorescence images were scored blindly for GFP expression in pollen and pollen tubes. Each GFP-expressing pollen grain or pollen tube was counted as one transformation event. Corresponding transmitted light images were used to obtain total pollen counts. Transformation efficiencies were calculated as a sum of transformation events over the total number of (ungerminated pollen grains plus germinated pollen tubes) counted in each treatment. Fisher's exact test was used to calculate p-values relative to the positive control for each individual trial. The experiment was conducted on three different days.

**Plasmid construct**

pUC19-260Zm13::GFP - harbors the -260 to +61 bp region of the maize *Zm13* gene (GRMZM2G317406) promoter upstream of the mGFP4 reporter gene [6] in pUC19. Gift of S. Li and E. Lord, UC Riverside.

**Pollen germination medium (PGM) [7]**

1 X PGM contains the following:

10% sucrose

0.0005% $H_3BO_3$

10 mM $CaCl_2$

0.05 mM $KH_2PO_4$

6% PEG 4000

After adding all components, PGM is heated to 70 °C for 10 min on stirring heated plate and filter sterilized.
